# Amyloid duration is associated with preclinical cognitive decline and tau PET

**DOI:** 10.1101/778415

**Authors:** Rebecca L. Koscik, Tobey J. Betthauser, Erin M. Jonaitis, Samantha L. Allison, Lindsay R. Clark, Bruce P. Hermann, Karly A. Cody, Jonathan W. Engle, Todd E. Barnhart, Charles K. Stone, Nathaniel A. Chin, Cynthia M. Carlsson, Sanjay Asthana, Bradley T. Christian, Sterling C. Johnson

## Abstract

**INTRODUCTION:** This study applies a novel algorithm to longitudinal amyloid positron emission tomography (PET) imaging to identify age-heterogeneous amyloid trajectory groups, estimate the age and duration (chronicity) of amyloid positivity, and investigate chronicity in relation to cognitive decline and tau burden.

**METHODS:** Cognitively unimpaired participants (n=257) underwent 1-4 amyloid PET scans. Group-based trajectory modeling was applied to participants with longitudinal scans (n=171) to identify and model amyloid trajectory groups, which were combined with Bayes’ theorem to estimate age and chronicity of amyloid positivity. Relationships between chronicity, cognition, clinical progression and tau PET (MK-6240) were investigated using regression models.

**RESULTS:** Chronicity explained more heterogeneity in amyloid binding than age and binary amyloid status. Chronicity was associated with faster cognitive decline, increased risk of abnormal cognition, and higher entorhinal tau.

**DISCUSSION:** Amyloid chronicity provides unique information about cognitive decline and neurofibrillary tangle development and may be useful to investigate preclinical AD.

## 1. INTRODUCTION

Alzheimer’s disease (AD) is characterized by beta-amyloid plaques and neurofibrillary tau tangles that accrue over time, leading to neurodegeneration and progressive cognitive and functional decline. Positron emission tomography (PET) biomarkers enable *in vivo* detection of pathophysiologic beta-amyloid and tau, and as hypothesized by Jack and colleagues^1,2^, these AD biomarkers follow nonlinear longitudinal patterns where detectable pathologic beta-amyloid accrues first^3,4^, perhaps twenty or more years prior to clinically detectable cognitive impairment^5,6^. The 2018 Research Framework for AD^7^ proposes that amyloid PET may be used to ascertain amyloid status (i.e. amyloid positive or negative; A+/−). This dichotomization is heuristically useful and multiple studies have shown that A+ cognitively unimpaired individuals exhibit greater cognitive decline over time than A−^8–10^, with greater cognitive decline for people who exhibit both elevated pathologic amyloid and tau^11,12^. However, among individuals who are accumulating amyloid, there is considerable heterogeneity in the magnitude and age of onset of amyloid accumulation with respect to age^13–15^. A method for elucidating such heterogeneity in amyloid accumulating cases may improve prediction models of the temporal biomarker cascade and cognitive decline.

As observed by Jack and colleagues in their seminal paper^1^, there is little known about inter-individual differences in middle-age beta-amyloid accumulation when individuals are transitioning from undetectable (A−) to detectable (A+) amounts of beta-amyloid. The theoretical sigmoidal model of beta-amyloid accumulation^1^ suggests that individuals have relatively slow accumulation initially, followed by faster accumulation as the disease progresses. Several approaches have examined ways to empirically assess the trajectory of beta-amyloid biomarkers in AD^6,13,16,17^. These approaches often attempt to align persons within a disease state to mitigate the biomarker heterogeneity with respect to age. A method that combines beta-amyloid magnitude and amyloid measurement age to estimate the age of biomarker onset (i.e. A+) could be useful in such scenarios since it would allow realignment of the time axis to describe the duration, or chronicity, of A+ relative to that person’s age at any given procedure. Group-based trajectory modeling (GBTM) is used to describe the developmental course(s) a phenomenon might follow over time^18,19^, and is well-suited for characterizing potential sub-distributions of PET measured beta-amyloid pathology accumulation patterns with respect to age. Modeling these sub-distributions in a sample containing amyloid convertors across the age spectrum may allow for more accurate estimation of the age at which persons become A+.

Using data from the Wisconsin Registry for Alzheimer’s Prevention (WRAP) study we investigated the following aims. First, we used longitudinal Pittsburgh compound B (PiB) to identify and characterize beta-amyloid trajectory groups of non-demented healthy middle-aged participants. Second, we examined whether trajectory group membership could be reliably obtained from only one PiB scan. We next utilized this trajectory group information to estimate the age of A+ (i.e., PiB positivity, [PIB(+)]) onset, and thereby the chronicity of A+ (i.e., time between estimated A+ onset and age at a given assessment). Third, we characterized mathematically the shape of the amyloid accumulation curve observed in this sample. Fourth, we investigated whether A+ chronicity was associated with cognitive decline and with tau tangles. Tau tangles were assessed using [^18^F]MK-6240, a novel PET radioligand with a high affinity for neurofibrillary tangles and minimal off-target binding in the brain ^20,21^.

## 2. METHODS

### 2.1 Sample

The sample included 257 WRAP participants who were cognitively unimpaired at baseline and completed at least one PiB PET scan as of June, 2019 (Table 1). WRAP is a longitudinal observational cohort study of late middle-aged and older adults, enriched for risk of AD by oversampling participants with a parental history of AD (73% parental AD history; see Johnson et al.,^20^). All study procedures were approved by the University of Wisconsin-Madison Institutional Review Board and are in concordance with the Helsinki declaration.

**Table 1.**
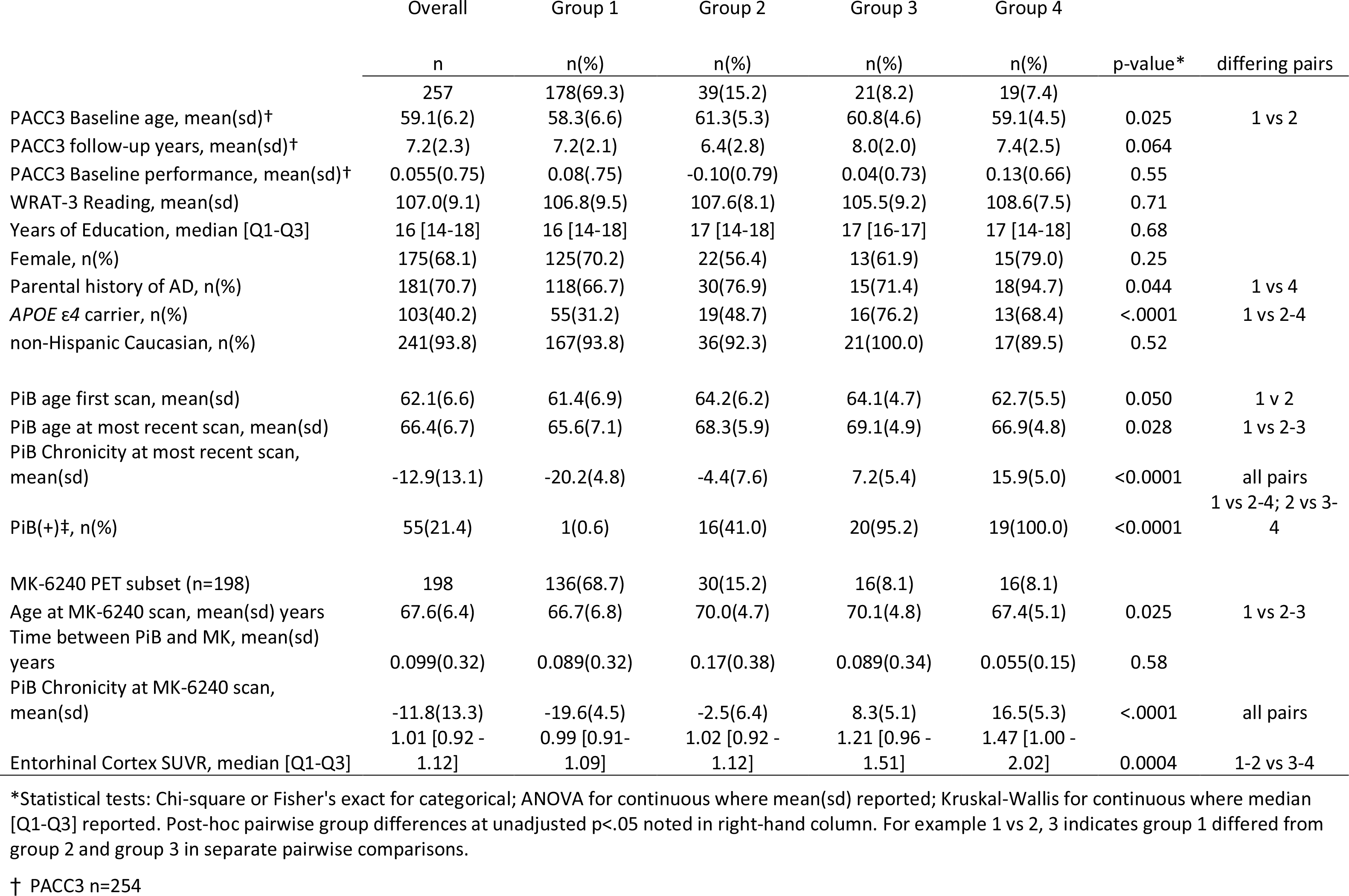
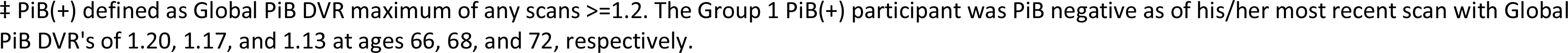
Sample characteristics, overall and by PiB Trajectory Group

### 2.2 Cognitive Assessment

WRAP participants complete cognitive assessments at baseline, and approximately every two years thereafter. Longitudinal cognitive performance was assessed using a preclinical Alzheimer cognitive composite score (PACC-3)^10,21^, derived from Rey Auditory Verbal Learning (RAVLT; Trials 1-5)^22^, Logical Memory II^23^, and Digit Symbol Substitution^24^.

### 2.3 Neuroimaging

All participants underwent T1-weighted magnetic resonance (MR) imaging, and [^11^C]PIB ([^11^C]6-OH-BTA-1)^25^. Amyloid burden was assessed as a global cortical average PIB distribution volume ratio (DVR)^26^ and a threshold of DVR ≥ 1.2^27^ to ascertain PIB(+); 198 also underwent [F-18]MK-6240 (6-(Fluoro-^18^F)-3-(1H-pyrrolo[2,3-c]pyridine-1-yl)isoquinolin-5-amine)) PET imaging^28^. Radioligand synthesis, and PET and MRI acquisition, processing, and analysis methods are described previously ^29,30^ and in *supplemental materials*.

### 2.4 Statistical Methods

Statistical analyses were conducted in SAS and R. Sample characteristics were compared across groups of interest (e.g., PiB trajectory groups) using tests appropriate for the distribution of the data.

#### 2.4.1 Aim 1

We used GBTM on the 171 participants with 2-4 PIB scans to identify PiB trajectory groups. GBTM is a special case of pattern mixture modeling in which individuals are classified into groups on the basis of longitudinal data^18,31,32^. Models are fit iteratively by adding and removing groups based on the Bayesian Information Criteria (BIC) fit statistics^18,31^. We modeled trajectories using up to a cubic polynomial, selecting the best parameterization based on BIC fit and reasonableness of the results. For example, if two functions had similar BIC for a group, the model that was more biologically probable was selected (i.e. accumulating groups were not allowed to estimate PiB DVR values less than the non-accumulating group).

#### 2.4.2 Aim 2

Using the GBTM functions and Bayes’ theorem (equation 1, Table 2) to estimate the probability of group membership in group “j” for each participant’s most recent scan; two re-weightings of the Bayes’ probabilities were applied to up-weight the probability of Group 1 or 2 for low Global PiB DVR values in age ranges where the trajectory functions were parallel and close (additional details described in supplement). Trajectory group membership was assigned as the group with maximum probability. Agreement between GBTM and Bayes’ theorem derived trajectory group assignments was examined using Kappa statisics^33^.

**Table 2:**
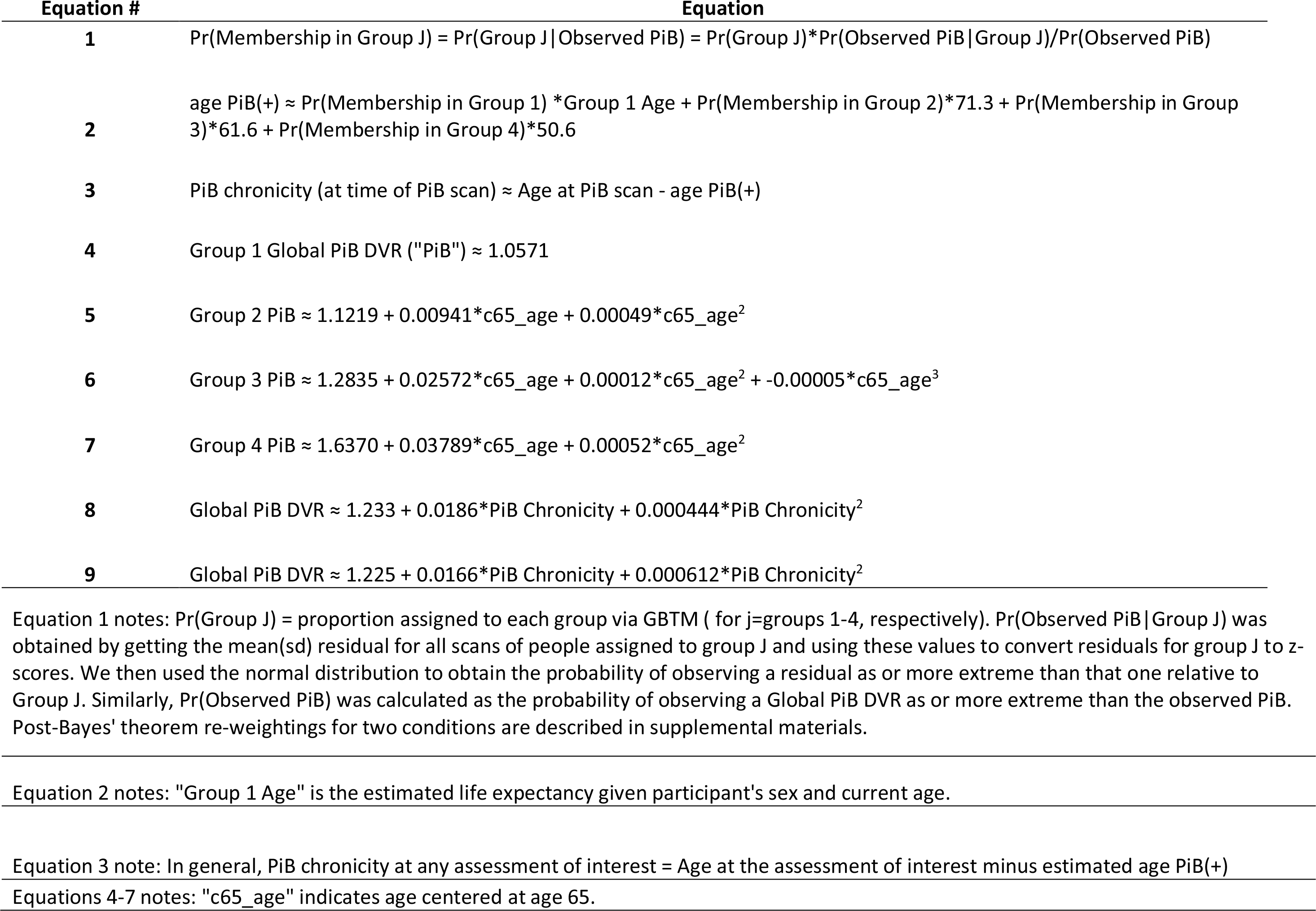
Equations.

#### 2.4.3 Aim 3

After observing strong agreement between PiB trajectory group assignment methods, the Bayes’ theorem approach was applied to all 257 participants to ascertain the probability of group membership and group assignment based on their Global PIB DVR at their most recent scan. A+ age was then estimated for each participant using a probability weighted average of the A+ ages of the trajectory groups (Table 2, equation 2). Amyloid chronicity was then calculated for each PET scan as the age at scan minus the estimated A+ age (Table 2, equation 3). By this convention, positive chronicity indicates the estimated duration of PIB(+) whereas negative chronicity indicates the person was PIB(−) at the time of the scan. Global PiB DVR was then modeled as a function of amyloid chronicity (including linear and quadratic chronicity terms). This function was used to estimate the time duration from the 10^th^ and 90^th^ PiB DVR centiles of the accumulating groups to the PiB(+) threshold to enable comparisons of amyloidosis duration with this method and sample to other studies.

#### 2.4.4 Aim 4

We used linear mixed effects (LME) models to examine whether amyloid chronicity at baseline PACC-3 modified longitudinal PACC-3 scores (random intercept and age-related slope; unstructured covariance; n=254 after excluding one participant with multiple sclerosis and two missing PACC-3 scores). Fits of the base model (covariates of sex, WRAT3, practice, age, age^2^) were compared with a model that included amyloid chronicity and its interaction with age and age^2^. After observing better model fit (lower corrected Akaike Information Criteria (AICc) statistics ^34^) for the latter model and significant interactions, we depicted the effects of amyloid chronicity on cognitive trajectories by plotting age trajectories for amyloid chronicity values that represented mean chronicity at PACC-3 baseline in each of the four PiB trajectory groups.

In secondary analyses, we used logistic regression to examine whether concurrent amyloid chronicity and age were associated with increased risk of abnormal cognition at most recent visit using three definitions of abnormal (progression to clinical impairment, abnormal relative to internal cross-sectional norms, and abnormal relative to longitudinal norms; see *supplemental materials*).

We used regression to compare age and amyloid chronicity at MK-6240 scan, PIB(+/−) status, and PIB DVR as predictors of entorhinal cortex MK-6240 SUVR. In separate models for each continuous predictor (age and amyloid chronicity), we began with cubic polynomial terms with the plan of sequentially removing non-significant highest order terms. To estimate how the PiB trajectory groups differed in terms of increase in MK-6240 per year of amyloid chronicity, we also used output from a model including a PiB trajectory group*chronicity interaction.

Sensitivity analyses were performed for all outcomes, substituting PiB(+/−) status and PIB DVR for amyloid chronicity and comparing AICc model fit statistics across otherwise identical models and consider |∆AICc| values <2 to represent comparable models^34^.

## 3. RESULTS

### 3.1 Aim 1

In the subset used for GBTM (n=171), mean(sd) age at first scan was 61.1(6.1) [range 46.9 – 78.9] with mean(sd)=5.8(2.0) years between first and last scan. Thirty-seven(21.6%) were PIB(+) for at least one scan, 114(66.7%) were female, 162(94.7%) non-Hispanic Caucasian, 70(40.9%) *APOE*4 carriers, and 124(72.5%) had a parental history of AD-dementia.

GBTM identified four age-defined PIB trajectory groups. Mean and median probabilities of group membership exceeded 70% in each group (indicating support for the four group solution). GBTM assigned 125(73.1%) to a non-accumulating group (Group 1), 21(12.3%) to a latest accumulating group (Group 2), 14(8.2%) to the middle accumulating group (Group 3), and 11(6.4%) to the earliest accumulating group (functions defining Groups 1-4 are Equations 4-7, respectively, in Table 2). Group functions and observed data are shown in Figure 1a.

**Figure 1.**
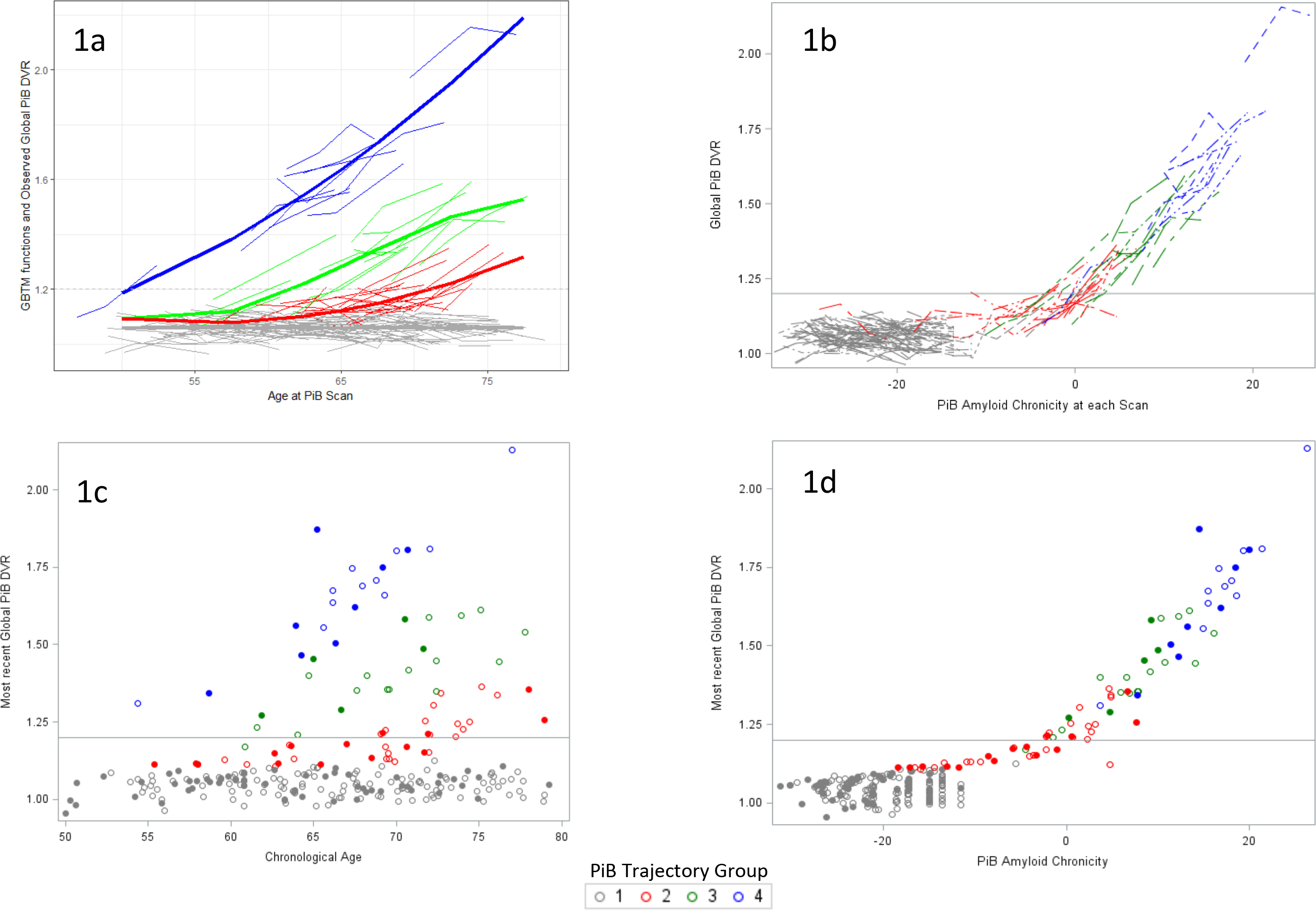
For all panels, gray indicates Group 1 (non-accumulators), red=Group 2, green=Group 3, blue=Group 4. Horizontal line indicates PiB(+) threshold (Global PiB DVR=1.2). a) Spaghetti plot of individual trajectories in set of 171 used in GBTM (thin lines) with 4 group functions identified by GBTM superimposed on the figure (thick lines; equations 4-7 in Table 2). b) Spaghetti plot of individual trajectories in set of 171 realigned vs amyloid chronicity. c) Scatter plot of most recent Global PiB DVR vs chronological age in expanded set (171 original = circles, 86 new people = dots; colors indicate trajectory group). d) Scatter plot of most recent Global PiB DVR vs amyloid chronicity in expanded set (coding same as above).

Solving equations 5-7 yielded estimated ages of PiB(+) of 71.3, 61.6, and 50.6 years for groups 2-4, respectively. Group 1 indicated an intercept only model that was below the PIB(+) threshold. Therefore, we estimated age of PiB(+) as age at PET scan plus life expectancy from a gender-specific life expectancy table (https://www.dhs.wisconsin.gov/stats/life-expectancy.htm; accessed 7/23/2019); this resulted in mean(sd) estimated age PiB(+) for group 1=88.0(2.2).

### 3.2 Aim 2

We observed strong agreement between PiB trajectory group assignment using GBTM (longitudinal scans) vs using trajectory group functions and Bayes’ theorem (only most recent scan in the GBTM set). Specifically, 160/171 group assignments agreed (93.6% agreement; Simple Kappa statistic=0.86, 95% CI=0.78-0.94) with perfect agreement in groups 3 and 4. GBTM and Bayes’ theorem derived group membership probabilities were highly correlated (Spearman, 0.87 for 160 concordant cases, 0.87 including 11 discrepancies; see supplement for details of discrepant cases). For all subsequent analyses, group membership based on Bayes’ theorem and most recent PIB scan is used.

### 3.3 Aim 3

Using the Bayes’ theorem approach we obtained group membership probabilities for all 257 participants, including 86 not included in GBTM modeling, which was used to assign PIB trajectory group membership (sample characteristics in Table 1). Equations 2 and 3 (Table 2) were then used to estimate amyloid chronicity for all participants based on their last PIB scan.

The four trajectory groups did not differ in terms of baseline PACC-3 performance, WRAT-III reading, years of education, race, or sex, but did differ in amyloid chronicity at most recent scan, parental history of AD and *APOE4* carriage. Follow-up pairwise comparisons among trajectory groups showed more *APOE4* carriage in each of the accumulating groups compared to group 1 and more parental history of AD in group 4 (the earliest accumulating group) compared to group 1.

Plots depicting PiB DVR vs chronicity and chronological age are shown in Figure 1 (1b longitudinal plots for the GBTM subset; 1c and 1d plotted cross-sectionally using most recent PIB). Chronicity and most recent PIB DVR were highly correlated (Pearson *r*=0.895) with a quadratic model indicating a good fit for chronicity predicting PIB DVR (R^2^=0.945 for all 257 participants, R^2^=0.931 including only groups 2-4, Table 2 equations 8 and 9, respectively). Using equation 9, we estimated it would take 10.0 years to go from PiB DVR=1.12 (10^th^ centile of accumulating groups) to PiB(+), and another 17.7 years to reach PIB DVR=1.71 (90^th^ centile of accumulating groups).

### 3.4 Aim 4

#### 3.4.1 Cognition

LME models of longitudinal PACC-3 showed better fit after adding the amyloid chronicity terms to the model including covariates and age terms (∆AICc decrease =−25.2). Interaction effects are depicted in Figure 2 for values representing mean amyloid chronicity at baseline PACC-3 in each of the PiB trajectory groups. In sensitivity analyses, substituting PiB(+/−) status for amyloid chronicity also resulted in better fit (∆AICc =−12.8) relative to base model, but not as good a fit as using chronicity (∆AICc =−12.5 model with amyloid chronicity AICc minus model with PiB(+) AICc).

**Figure 2.**
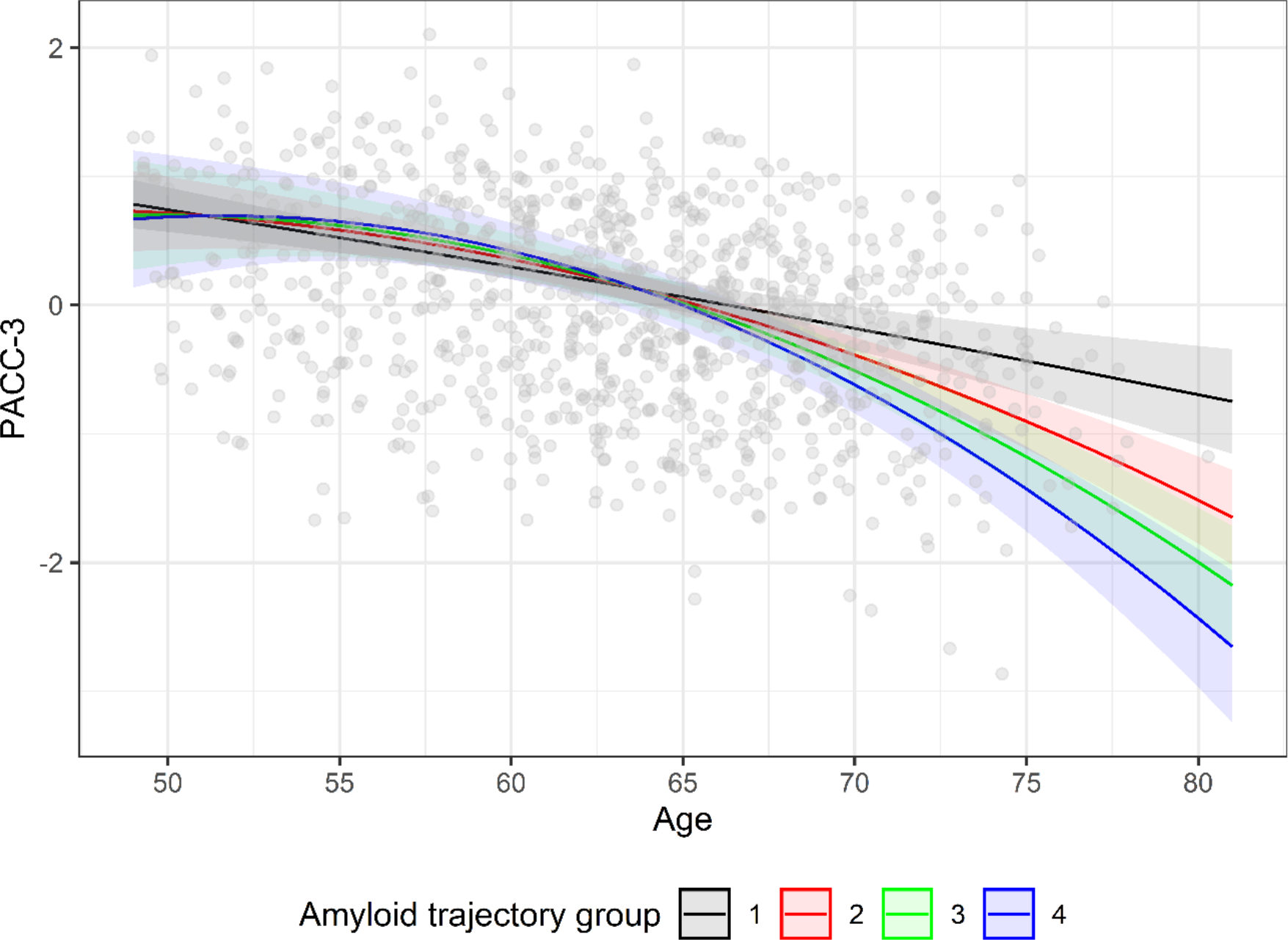
Interaction plot from LME of PACC-3. Lines depict age trajectories for PiB chronicities of −28, −11, −1 and 8 (these values are the mean PiB chronicity at baseline PACC-3 of Groups 1-4, respectively). Dots indicate observed PACC-3 values. Predicted PACC-3 ≈ −0.3658 + −0.4304*Male + 0.0291*c100_WRAT3 + 0.1142*Practice +−0.09149*c65_age + −0.00182*PiB chronicity + −0.00305*c65_age^2^+ −0.00156*c65_age*PiB chronicity + −0.00010* age^2^*PiB chronicity (and random person-level intercepts and age slopes); c100_WRAT3 indicates WRAT3 reading standard score, centered at value of 100 and c65_age indicates age centered at 65.

Logistic regression showed a consistent pattern of statistically significant risk of abnormal cognitive status associated with amyloid chronicity but not age where abnormal cognitive status was defined relative to clinical criteria and internal norms^35^. Odds ratios and CIs for age and amyloid chronicity are shown in Figure 3 for each of these outcomes. Compared to those who did not progress to MCI/AD (n=238), those who progressed (n=16) were on average 4.8(6.2) years older at their most recent cognitive assessment, but were estimated to be amyloid positive for 15.5(12.5) years longer. Similarly, age and amyloid chronicity were 0.93(6.2) and 11.2(12.6) years higher in those below (n=27) vs above (n=218) the cross-sectional internal norms cut-off; and age and PiB chronicity were 2.8(6.2) and 11.2(12.6) years higher in those below (n=28) vs above (n=217) the longitudinal internal norm cut-off. Sensitivity analyses substituting PIB(+/−) status for chronicity indicated worse fit statistics for all cognitive outcomes (∆AICc range = −6.2 to −8.4) compared to chronicity. Substituting last PIB DVR for chronicity indicated chronicity was a better fit for predicting MCI/AD and abnormal cross-sectional norms (∆AICc’s= −3.1 and −3.3, respectively) but not for abnormal longitudinal norms (∆AICc=2.8).

**Figure 3.**
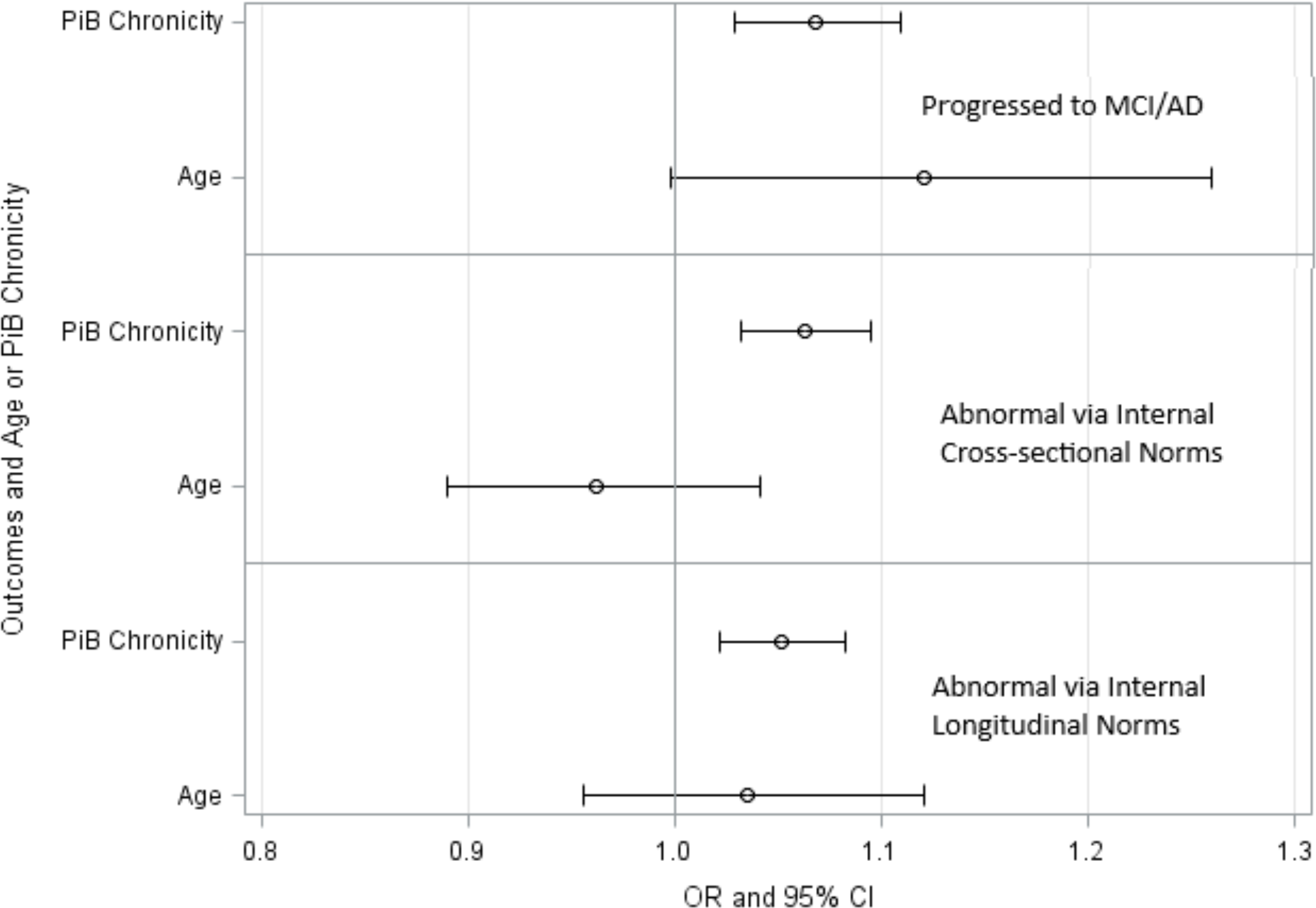
Forest plots of OR’s with various outcomes indicating abnormal at last cognitive assessment. The top pair of variables indicates odds ratios (OR’s) and their 95% CI’s for predicting progression from Cognitively Unimpaired to MCI or Dementia (cognitive statuses determined by consensus conference as described in^20^). The middle pair of variables show OR’s and CI’s for predicting an abnormal PACC-3 score at the most recent cognitive assessment according to internal demographically-adjusted cross-sectional norms (i.e., <= 7^th^ centile, or ~1.5 SD or more below expected). The bottom pair of variables show OR’s and CI’s for predicting an abnormal change in PACC-3 score at the most recent cognitive assessment according internal longitudinal norms (i.e., <= 7^th^ longitudinal centile).

#### 3.4.2 Entorhinal Tau

One-hundred ninety-eight participants (77%) also underwent MK-6240 PET scans (mean(sd) of 0.10(0.32) years between last PiB and MK-6240 scans; mean(sd) age at MK-6240 scan=67.6(6.4)). Amyloid chronicity at time of MK-6240 differed between all PiB trajectory groups in a stepwise manner (Table 1). Mean entorhinal MK-6240 SUVR was near one for groups 1 and 2, increased stepwise for groups 3 and 4, and indicated significant group differences (p = 0.0004; groups 1&2 differed from groups 3&4). Only the linear age term was a significant predictor of entorhinal cortex MK-6240 SUVR; in a separate model, all terms in the cubic amyloid chronicity polynomial model were significant predictors of entorhinal cortex MK-6240 SUVR (Figure 4; ∆AICc chronicity – age models = −120.7). Including the trajectory group term and its interaction with chronicity in the model indicated that MK-6240 SUVR values were 1.1, 3, and 10.6 times higher for the amyloid accumulating groups (i.e. groups 2-4, respectively) compared to the non-accumulating group (i.e. group 1; interaction p-value ~.0001; ∆AICc chronicity polynomial – group*chronicity interaction model = −6.5).

**Figure 4.**
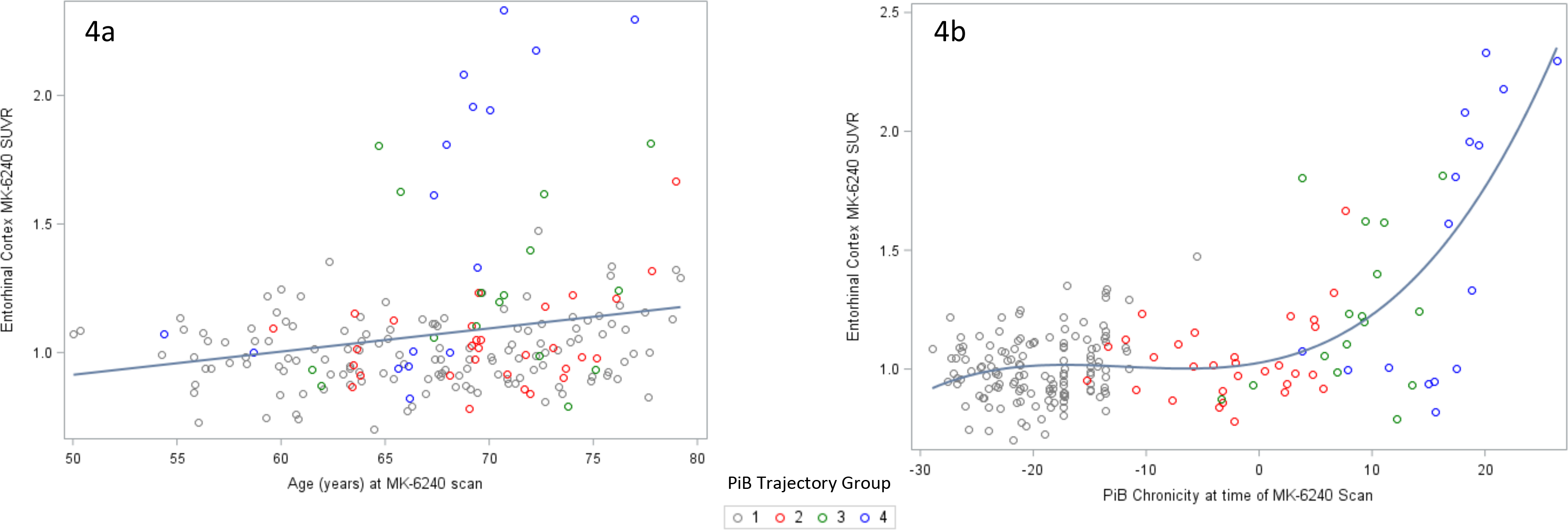
Entorhinal cortex SUVR: 4a) chronological age vs MK SUVR (model reduced sequentially from cubic polynomial to model including only linear age term); 4b) PiB chronicity vs MK SUVR (all three time terms in cubic polynomial were significant). Colors indicate PiB trajectory group (gray=non-accumulators, red=Group 2, green=Group 3, blue=Group 4).

Sensitivity analyses replacing chronicity with PiB(+/−) status indicated PiB(+/−) status was also a significant predictor of MK-6240 entorhinal cortex SUVR, although it was a weaker predictor than chronicity (Figure 4; ∆AICc chronicity – PiB(+) models=−81.5). Replacing amyloid chronicity with Global PiB DVR yielded models with similar fit (∆AICc chronicity – Global PiB DVR models = −1.3).

## 4. DISCUSSION

This work demonstrates a novel application of group-based trajectory modeling and Bayes’ theorem to identify amyloid trajectory groups with respect to chronological age and to estimate the age of amyloid positivity within groups and individuals, and thereby the time duration of exposure to pathologic beta-amyloid (i.e., amyloid chronicity). This methodology was applied to investigate differences between trajectory groups, and relationships between amyloid duration, chronological age, PET measured neurofibrillary tangles, and cognitive trajectories of initially cognitively unimpaired persons. Among the key findings was that the estimated duration of amyloid positivity (i.e. amyloid chronicity) maintained information about dichotomous amyloid status while simultaneously preserving information about the severity of amyloidosis. In addition, this approach provided insights regarding the heterogeneity of amyloid accumulation trajectories with respect to age that become homogeneous when reordering the time-axis to reflect the estimated duration of amyloidosis.

### 4.1 Beta-amyloid modeling and rates of accumulation

Previous studies have used several different approaches and cohorts to model amyloid trajectories with respect to age and amyloid biomarker levels^1,14,16,36,37^. These studies suggest that individuals in the AD continuum begin accumulating beta-amyloid at different ages, but experience similar rates of amyloid accumulation for a given level of amyloidosis. In agreement with these studies the GBTM results suggest there exist subgroups of amyloid accumulators that differ in the age of PET detectable amyloidosis onset, but the rates of amyloid accumulation with respect to time of amyloid onset are similar across individuals. This was demonstrated in this study by plotting the longitudinal amyloid data along the time axis realigned for duration of amyloid positivity and observing the similarity in slopes across subjects with similar durations of estimated amyloid positivity. The shape of this data was also consistent with the early portion of the beta-amyloid biomarker model of Jack and colleagues (Jack 2013). However, we did not observe evidence of a slowing of the accumulation rate, which was potentially due to the sample being younger and primarily asymptomatic and therefore earlier in the Alzheimer’s disease continuum compared to other studies. Further work adding new longitudinal cases will be needed to examine to what extent the homogeneity in this curve is maintained in the presence of new longitudinal scans not used to fit the initial trajectory functions.

### 4.2 Chronology of beta-amyloid relative to tau and cognitive decline

Understanding the chronology of AD biomarkers and their prognostic value is important for contextualizing studies relating AD biomarkers to symptomology and other disease outcomes, and for clinical trial design. Previous studies have proposed different methods for obtaining metrics reflective of disease state^1,3,16,36–38^. In contrast to those methods, the approach in this study uses fewer model parameters and less complex model functions. Further, the relative timing between chronological events is not affected by the positivity threshold, the output (time duration) is easily interpretable, and estimates can be obtained from a single cross-sectional PET scan once the group model is trained. A criticism of this approach is that amyloid chronicity is highly correlated with DVR estimates (r=0.90). However, by recasting the magnitude of amyloid elevation along the time dimension, a number of important questions become contextualized and more readily addressable, including determining the effect of putative risk/resilience factors (e.g. hypertension, physical activity) on the onset of AD amyloidosis and or AD-associated cognitive decline.

As initial proof-of-concept, the relationships between the estimated amyloid chronicity (i.e. duration of amyloid positivity) and markers of tau pathophysiology and cognition were investigated in a late middle-aged, mostly asymptomatic sample. These analyses suggested that those who were amyloid positive for a greater length of time at cognitive baseline exhibited faster rates of cognitive decline during the 7.2 years of cognitive follow-up, and that persons who developed clinical levels of cognitive impairment were estimated to have been A+ for a mean of 15 years longer than those who did not convert to clinical impairment. These intuitive results may partly explain why some studies demonstrate a relationship between A+ status and cognition during the preclinical stage, whereas others do not.^8,39–41^

Similar to relationships with cognition, models including chronicity to explain entorhinal tau tangles improved model fits compared to models with age alone or dichotomous amyloid status. Further, these results suggested a time-lapsed relationship wherein tangles were detectable several years after the pathologic beta-amyloid. In agreement with previous studies, these results support the hypotheses that the level of amyloid tracer binding is reflective of the cumulative process of amyloidosis, and that markers of AD pathophysiology and cognition follow a temporal hierarchy. In addition, these results support previous findings that suggest age is a risk factor for pathophysiology and cognitive decline in AD, but age itself is not a robust predictor of amyloidosis, entorhinal tangles and thereby Alzheimer’s disease state.

The major contribution of this “proof of concept” study is that the trajectory of amyloid accumulation, age of amyloid positivity, and its chronicity can be estimated and used to describe the disease course of amyloidosis. Study limitations include the following. WRAP is a volunteer cohort with over-sampling of participants with a parental history of AD; this results in higher AD-risk characteristics than in the general population. Additionally, the sample is younger and more cognitively intact compared to other longitudinal studies of amyloid accumulation; thus, it is unlikely that our exact equations and parameters will generalize to other radiotracers and study samples. As such, replication of this method in different cohorts is needed to determine to what extent this approach is generalizable. If replicable, estimates of amyloid onset and chronicity, such as described here, should be examined in other research contexts to better understand the impact of treatments, preventative measures, and resilience factors during the preclinical phase of AD.

## ABBREVIATIONS

AD: Alzheimer’s disease
PET: positron emission tomography
A+/−: amyloid positive/negative
GBTM: group-based trajectory modeling
WRAP: Wisconsin Registry for Alzheimer’s Prevention
PIB: Pittsburgh Compound B
PIB(+/−): Pittsburgh Compound B positive/negative
PACC-3: three-test preclinical Alzheimer cognitive composite
RAVLT: Rey Auditory Verbal Learning
MR: magnetic resonance
DVR: distribution volume ratio
BIC: Bayesian Information Criteria
WRAT3: Wide Range Achievement Test 3
AICc: Akaike Information Criteria
sd: standard deviation
*APOE4*: apolipoprotein ε4
LME: linear mixed effects

## ACKNOWLEDGMENTS

Funding: This work was supported by the National Institutes of Health [R01 AG021155 y6-y13, R01 AG027161 y1-y11, P50 AG033514 y1-y11, U54 HD090256 y1-y4], the Wisconsin Alzheimer’s Institute Lou Holland Fund and the Alzheimer’s Association [AARF-19-614533, 2019]. These funding sources did not contribute to study design, analysis, interpretation or writing of this work. We would also like to thank all participants and their families, and the many study teams at the University of Wisconsin-Madison that make this work possible.

## References

1. Jack JrCR, Knopman DS, Jagust WJ, et al. Tracking pathophysiological processes in Alzheimer’s disease: an updated hypothetical model of dynamic biomarkers. Lancet Neurol. 2013;12(2):207–216.

2. Jack JrCR, Knopman DS, Jagust WJ, et al. Hypothetical model of dynamic biomarkers of the Alzheimer’s pathological cascade. Lancet Neurol. 2010;9(1):119–128.

3. Bateman RJ, Xiong C, Benzinger TL, et al. Clinical and biomarker changes in dominantly inherited Alzheimer’s disease. N Engl J Med. 2012;367(9):795–804.

4. Rowe CC, Ellis KA, Rimajova M, et al. Amyloid imaging results from the Australian Imaging, Biomarkers and Lifestyle (AIBL) study of aging. Neurobiol Aging. 2010;31(8):1275–1283.

5. Jack JrCR, Lowe VJ, Weigand SD, et al. Serial PIB and MRI in normal, mild cognitive impairment and Alzheimer’s disease: implications for sequence of pathological events in Alzheimer’s disease. Brain. 2009;132(5):1355–1365.

6. Villemagne VL, Burnham S, Bourgeat P, et al. Amyloid β deposition, neurodegeneration, and cognitive decline in sporadic Alzheimer’s disease: a prospective cohort study. Lancet Neurol. 2013;12(4):357–367.

7. Jack CR, Bennett DA, Blennow K, et al. NIA-AA Research Framework: Toward a biological definition of Alzheimer’s disease. Alzheimers Dement J Alzheimers Assoc. 2018;14(4):535–562.

8. Clark LR, Berman SE, Norton D, et al. Age-accelerated cognitive decline in asymptomatic adults with CSF β-amyloid. Neurology. 2018;90(15):e1306–e1315.

9. Clark LR, Racine AM, Koscik RL, et al. Beta-amyloid and cognitive decline in late middle age: findings from the Wisconsin Registry for Alzheimer’s Prevention study. Alzheimers Dement. 2016;12(7):805–814.

10. Donohue MC, Sperling RA, Petersen R, Sun C-K, Weiner MW, Aisen PS. Association between elevated brain amyloid and subsequent cognitive decline among cognitively normal persons. Jama. 2017;317(22):2305–2316.

11. Desikan RS, McEvoy LK, Thompson WK, et al. Amyloid-β–associated clinical decline occurs only in the presence of elevated p-tau. Arch Neurol. 2012;69(6):709–713.

12. Sperling RA, Mormino EC, Schultz AP, et al. The impact of amyloid-beta and tau on prospective cognitive decline in older individuals. Ann Neurol. 2019;85(2):181–193.

13. Bilgel M, Prince JL, Wong DF, Resnick SM, Jedynak BM. A multivariate nonlinear mixed effects model for longitudinal image analysis: Application to amyloid imaging. Neuroimage. 2016;134:658–670.

14. Villemagne VL, Burnham S, Bourgeat P, et al. Amyloid β deposition, neurodegeneration, and cognitive decline in sporadic Alzheimer’s disease: a prospective cohort study. Lancet Neurol. 2013;12(4):357–367.

15. Jack CR, Wiste HJ, Lesnick TG, et al. Brain β-amyloid load approaches a plateau. Neurology. 2013;80(10):890–896.

16. Insel PS, Ossenkoppele R, Gessert D, et al. Time to amyloid positivity and preclinical changes in brain metabolism, atrophy, and cognition: evidence for emerging amyloid pathology in Alzheimer’s disease. Front Neurosci. 2017;11:281.

17. Whittington A, Sharp DJ, Gunn RN, Initiative ADN. Spatiotemporal distribution of β-amyloid in Alzheimer disease is the result of heterogeneous regional carrying capacities. J Nucl Med. 2018;59(5):822–827.

18. Jones BL, Nagin DS, Roeder K. A SAS procedure based on mixture models for estimating developmental trajectories. Sociol Methods Res. 2001;29(3):374–393.

19. Nagin DS. Analyzing developmental trajectories: a semiparametric, group-based approach. Psychol Methods. 1999;4(2):139.

20. Johnson SC, Koscik RL, Jonaitis EM, et al. The Wisconsin Registry for Alzheimer’s Prevention: A review of findings and current directions. Alzheimers Dement Diagn Assess Dis Monit. 2018;10:130–142. doi:10.1016/j.dadm.2017.11.007

21. Jonaitis EM, Koscik RL, Clark LR, et al. Measuring longitudinal cognition: Individual tests versus composites. Alzheimers Dement Diagn Assess Dis Monit. 2019;11:74–84.

22. Rey A. L’examen psychologique dans les cas d’encéphalopathie traumatique.(Les problems.). Arch Psychol. 1941.

23. Wechsler D. WMS-R: Wechsler Memory Scale-Revised. Psychological Corporation; 1987.

24. Wechsler D. WAIS-III: Wechsler Adult Intelligence Scale. Psychological Corporation; 1997.

25. Klunk WE, Lopresti BJ, Ikonomovic MD, et al. Binding of the positron emission tomography tracer Pittsburgh compound-B reflects the amount of amyloid-β in Alzheimer’s disease brain but not in transgenic mouse brain. J Neurosci. 2005;25(46):10598–10606.

26. Sprecher KE, Bendlin BB, Racine AM, et al. Amyloid burden is associated with self-reported sleep in nondemented late middle-aged adults. Neurobiol Aging. 2015;36(9):2568–2576.

27. Racine AM, Clark LR, Berman SE, et al. Associations between performance on an abbreviated CogState battery, other measures of cognitive function, and biomarkers in people at risk for Alzheimer’s disease. J Alzheimers Dis. 2016;54(4):1395–1408.

28. Hostetler ED, Walji AM, Zeng Z, et al. Preclinical characterization of 18F-MK-6240, a promising PET tracer for in vivo quantification of human neurofibrillary tangles. J Nucl Med. 2016;57(10):1599–1606.

29. Betthauser TJ, Cody KA, Zammit MD, et al. In Vivo Characterization and Quantification of Neurofibrillary Tau PET Radioligand 18F-MK-6240 in Humans from Alzheimer Disease Dementia to Young Controls. J Nucl Med. 2019;60(1):93–99.

30. Johnson SC, Christian BT, Okonkwo OC, et al. Amyloid burden and neural function in people at risk for Alzheimer’s disease. Neurobiol Aging. 2014;35(3):576–584.

31. Jones BL, Nagin DS. Advances in group-based trajectory modeling and an SAS procedure for estimating them. Sociol Methods Res. 2007;35(4):542–571.

32. Nagin DS, Odgers CL. Group-based trajectory modeling in clinical research. Annu Rev Clin Psychol. 2010;6:109–138.

33. McHugh ML. Interrater reliability: the kappa statistic. Biochem Medica Biochem Medica. 2012;22(3):276–282.

34. Burnham KP, Anderson DR, Huyvaert KP. AIC model selection and multimodel inference in behavioral ecology: some background, observations, and comparisons. Behav Ecol Sociobiol. 2011;65(1):23–35.

35. Koscik RL, Jonaitis EM, Clark LR, et al. Longitudinal standards for mid-life cognitive performance: Identifying abnormal within-person changes in the Wisconsin Registry for Alzheimer’s Prevention. J Int Neuropsychol Soc. 2019;25(1):1–14.

36. Whittington A, Sharp D, Gunn R. Spatiotemporal distribution of β-amyloid in Alzheimer’s Disease results from heterogeneous regional carrying capacities. J Nucl Med. 2016;57(supplement 2):346–346.

37. Bilgel M, An Y, Zhou Y, et al. Individual estimates of age at detectable amyloid onset for risk factor assessment. Alzheimers Dement. 2016;12(4):373–379.

38. Jedynak BM, Lang A, Liu B, et al. A computational neurodegenerative disease progression score: method and results with the Alzheimer’s disease Neuroimaging Initiative cohort. Neuroimage. 2012;63(3):1478–1486.

39. Aizenstein HJ, Nebes RD, Saxton JA, et al. Frequent amyloid deposition without significant cognitive impairment among the elderly. Arch Neurol. 2008;65(11):1509–1517.

40. Mormino EC, Kluth JT, Madison CM, et al. Episodic memory loss is related to hippocampal-mediated β-amyloid deposition in elderly subjects. Brain. 2008;132(5):1310–1323.

41. Pike KE, Savage G, Villemagne VL, et al. β-amyloid imaging and memory in non-demented individuals: evidence for preclinical Alzheimer’s disease. Brain. 2007;130(11):2837–2844.

