## Supplemental materials for "Amyloid duration is associated with preclinical cognitive decline and tau PET"

**Methods**

**Additional details for section 2.3 Neuroimaging:**

**MRI**: T1-weighted brain volume scans were acquired on a 3T GE MR750 scanner with a 3D inversion recovery prepared fast spoiled gradient-echo sequence which was then bias corrected, tissue class segmented, and spatially normalized to MNI152 template space (SPM12). Automated anatomical labeling[1] and Harvard-Oxford[2] atlases were inverse warped to native T1 space and restricted to gray matter probabilities greater than 0.3. PET regions of interest (ROIs) were generated by smoothing binary masks by an isotropic 6 mm Gaussian kernel and keeping voxels greater than 0.7.

**PET:** All PET images were acquired on either a Siemens EXACT HR+ or a Siemens Biograph Horizon tomograph at the University of Wisconsin-Madison Waisman Center for Brain Imaging. Reconstructed PET time series were corrected from photon attenuation and scatter, radioactive decay, detector normalization, random events, and dead time.

**PET PiB:** Seventy-minute dynamic PiB data (5 x 2 min, 12 x 5 min frames) were acquired beginning with nominal bolus injection of 370 MBq [$^{11}$C]PiB. The PET time series was smoothed (3 mm isotropic Gaussian), interframe realigned, denoised (HYPR-LR[3]), and coregistered to T1-weighted MRI. The distribution volume ratio (DVR) was estimated using reference Logan graphical analysis[4] ($k_2' = 0.149$ min$^{-1}$, $t^* = 35$ min, cerebellar gray matter reference region) as a measure of beta-amyloid plaque burden. A global average PiB DVR was determined by taking the mean of 8-bilateral regions of interest from the AAL atlas[5] and a threshold of global PiB DVR ≥ 1.2[6] was used to ascertain PiB positivity [PiB(+)].

**PET MK-6240:** MK-6240 data were dynamically (4 x 5 min frames) acquired beginning 70-minutes after nominal 370 MBq MK-6240 injection. The MK-6240 time series were smoothed (6 mm isotropic Gaussian kernel), interframe realigned, summed, and coregistered to T1-weighted MRI.

Standard uptake value ratio (SUVR) was calculated using the inferior cerebellum as a reference region of non-displaceable binding. Regional SUVR values were extracted using the gray matter restricted Harvard-Oxford Atlas ROIs. The bilateral mean MK-6240 SUVR in the entorhinal cortex (anterior parahippocampal gyrus) was used as a measure of neurofibrillary tangle burden. This region was chosen because of its typical early involvement in AD.

**Additional details for section 2.4.2 Aim 2:**

As noted in the main text (Section 2.4.2), we used the four GBTM functions (equations 4-7, Table 2) and Bayes' theorem (equation 1, Table 2) to estimate the probability of group membership in group "j" for each participant's most recent scan. Group membership was preliminarily assigned as the group with maximum probability of group membership via Bayes' theorem. Visual inspection of PiB trajectory group assignment relative to trajectory functions, age, and Global PiB DVR identified potential for misclassification relative to Group 1 either at low PiB DVR values and the younger ages when the Group 1, 2 and 3 functions were close and relatively parallel with each other (see thick lines in main paper Figure 1a) or Global PiB DVR values that were below the PiB(+) threshold but well above the Group 1 mean for persons at the upper end of our observed age range. Two re-weightings of the Bayes' probabilities were applied to improve the accuracy of estimating age PiB(+) in these situations. First, for a Group 1 residual <0 (i.e., Global PiB DVR below non-accumulator group mean) and Bayes'-based PiB trajectory group-assignment in one of the accumulating groups (i.e., Groups 2-4), we multiplied the Group 1 Bayes' probability by 4 and the Group 2 Bayes' probability by 2 and then used these to obtain re-weighted probabilities of group membership. In the n=257, this resulted in reassigning one person who was ~age 50 at his/her scan and had a Global PiB DVR<1.0 from Group 3 via Bayes' theorem to Group 1 with the re-weighting.  Similarly, for those who were assigned to Group 1 via Bayes' probabilities but had Group 1 residual z-scores > 1.5 (i.e., had Global PiB DVR's that overlapped with those observed in Group 2), we multiplied Group 2 probabilities by 4 and obtain re-weighted probabilities of group membership. Group assignment changed from Group 1 to 2 for a participant ~age 69 with Global PiB DVR=1.12. Reweighting was done prior to characterizing the Global PiB DVR vs PiB Chronicity curve and prior to using PiB Chronicity as a predictor of other variables.

**Additional details for section 2.4.4**

As noted in the main text (Section 2.4.4) we used logistic regression to examine whether PiB chronicity was a stronger predictor of categorical cognitive outcomes than chronological age. Categorical outcomes included (i) progression to MCI or dementia as determined by independent consensus diagnosis conference [7]; (ii) abnormal WRAP-PACC-3 performance at last cognitive visit on unconditional (cross-sectional) demographically-adjusted internal norms; or (iii) abnormal WRAP-PACC-3 performance at last cognitive visit on conditional (longitudinal) demographically-adjusted internal norms. For (ii) and (iii), cross-sectional and longitudinal norms were developed for the WRAP-PACC-3 following the methods described in Koscik, Jonaitis et al.[8] with the following adjustments: the outcome was modeled using restricted regression quantiles[9,10], and for the longitudinal norms, baseline performance was operationalized as the first measurement only rather than the mean of the first two. We defined 'abnormal' for (ii) and (iii) using <= the 7$^{th}$ centile as this corresponds approximately to -1.5 sd below expectation relative to internal norms.

### Results

**Additional details for section 3.2 Aim 2 – PiB trajectory group: Discrepancies in GBTM vs Bayes' theorem assignment.**

Six of the 11 discrepant cases included participants who were assigned to group 1 via the GBTM program and group 2 via Bayes' theorem. For all six, their last PiB DVR was >= 1.85 standard deviations above the mean for Group 1 (range = 1.85 to 3.15 SD above the mean), suggesting that group 2 was a reasonable assignment based on the last scan. Three additional discrepancies included participants who were assigned to group 2 via the GBTM program and group 1 via Bayes' theorem at their last scan; one of these three had a PiB DVR that was .68 SD above the group 1 mean, suggesting group 1 was a reasonable assignment based on the single scan. This person also showed a decline from previous scans that suggested measurement error may have been a factor in the discrepancy. The other two cases, however, at PiB ~ 1.12 were > 2 sds above the group 1 mean, suggesting Group 2 was a more appropriate assignment based on most recent scan. Two other participants were assigned to group 2 via GBTM and group 3 via Bayes' theorem. Each of these participants showed large increases in PiB from prior scans suggesting that the group 3 assignment based on last scan was reasonable. Overall, this additional qualitative analyses of the discrepant cases suggested the Bayes' theorem approach was a reasonable way to use the four PiB trajectory functions to estimate probability of group membership and assign group membership using a single scan.
